# Spatio-temporal point processes as meta-models for population dynamics in heterogeneous landscapes

**DOI:** 10.1101/2021.06.04.447081

**Authors:** Patrizia Zamberletti, Julien Papaïx, Edith Gabriel, Thomas Opitz

## Abstract

Landscape heterogeneity affects population dynamics, which determine species persistence, diversity and interactions. These relationships can be accurately represented by advanced spatially-explicit models (SEMs) allowing for high levels of detail and precision. However, such approaches are characterised by high computational complexity, high amount of data and memory requirements, and spatio-temporal outputs may be difficult to analyse. A possibility to deal with this complexity is to aggregate outputs over time or space, but then interesting information may be masked and lost, such as local spatio-temporal relationships or patterns. An alternative solution is given by meta-models and meta-analysis, where simplified mathematical relationships are used to structure and summarise the complex transformations from inputs to outputs. Here, we propose an original approach to analyse SEM outputs. By developing a meta-modelling approach based on spatio-temporal point processes (STPPs), we characterise spatio-temporal population dynamics and landscape heterogeneity relationships in agricultural contexts. A landscape generator and a spatially-explicit population model simulate hierarchically the pest-predator dynamics of codling moth and ground beetles in apple orchards over heterogeneous agricultural landscapes. Spatio-temporally explicit outputs are simplified to marked point patterns of key events, such as local proliferation or introduction events. Then, we construct and estimate regression equations for multi-type STPPs composed of event occurrence intensity and magnitudes. Results provide local insights into spatio-temporal dynamics of pest-predator systems. We are able to differentiate the contributions of different driver categories (*i*.*e*., spatio-temporal, spatial, population dynamics). We highlight changes in the effects on occurrence intensity and magnitude when considering drivers at global or local scale. This approach leads to novel findings in agroecology where the organisation of cultivated fields and semi-natural elements are known to play a crucial role for pest regulation. It aids to formulate guidelines for biological control strategies at global and local scale.

## 1 Introduction

Community structure, population dynamics and species interactions within and between trophic levels are not limited within single plot’s borders but depend on the spatial context (*e*.*g*., patch size, spatial configuration, landscape composition, habitat connectivity; see Delaune et al. (2019)) and on ecological processes at different spatial scales (Pickett and Siriwardena, 2011). The key to understanding and predicting community structure and population distribution lies in the explication of the latent mechanisms and causes underlying observed patterns, which may emerge from the collective behaviour at smaller scale units or may be imposed by larger-scale constraints and the related temporal scale (Levin, 1992). Moreover, the influence of different spatial and temporal scales is closely related with species life-history traits, such as their ability to disperse, body size, competition, habitat specialisation, or trophic position (Rusch et al., 2010; O’Rourke et al., 2011). For example, foraging range and dispersal ability may determine the landscape elements that contribute to population dynamics and trophic interactions (Eber, 2001; Fahrig, 2001; Tscharntke and Brandl, 2004). Changes in spatial arrangement of habitats and composition could induce investment in the adaptation of dispersal-related traits (Tscharntke and Brandl, 2004).

Hence, dealing with ecological processes involves studying different spatial and temporal scales, since ecosystem patterns and processes cover various spatio-temporal ranges and may have multiple drivers acting across different extents (Fritsch et al., 2020). The characterisation of the spatial distribution of landscape features and individuals in response to such complex interplay of processes across scales belongs to the field of *landscape ecology*. To account for this complexity, the development of spatially explicit computer modelling and simulations are central for addressing theoretical questions. Many Spatially Explicit Model (SEM) types have been proposed, such as continuous-space reaction-diffusion partial differential equations (Roques, 2013), patch models (Hanski and Thomas, 1994), cellular automata neighborhood models (Hogeweg, 1988), or individual-based models (IBM, Grimm et al., 2005). DeAngelis and Yurek (2017) show the importance and the benefits of using SEMs compared to Spatially Implicit Models (SIMs) through different examples, including a savanna ecosystem. They find that the details and small-scale processes captured by SEMs are fundamental drivers for the ecosystem and its dynamics. SEMs can simulate the emergence of both small- and large-scale patterns from these processes and reveal deep details of dynamics such as predator–prey interactions and food web chains.

The development of advanced numerical models has greatly improved our ability to accurately describe complex dynamics incorporating fine-grain interactions over a large extent. However, as models aim to provide a realistic but simplified representation of reality, the spatio-temporal extent is often properly adapted by scaling decisions (Fritsch et al., 2020). In-model scaling methods give control over simplifications when building the model or allow us to incorporate and transfer relevant information across different scales. Scaling techniques may also be used before or after building the model, to define model parameters or analyse model outputs. In this work we focus on post-model scaling and propose a parsimonious approach to deal with the complexity of SEM outputs while keeping fine-scale information on the ecological dynamics. A solution to deal with this complexity could be the application of non-spatial analysis methods via spatial and temporal output aggregation (Gotelli, 2000; Webb, 2000; Fritsch et al., 2020). For example, Nathan et al. (2019) use spatially-explicit IBMs to study the hybridisation dynamics among species by describing their relationships across ecological scales, and then model outputs are integrated over space and time. In this case, however, all fine-scale information is lost, thus impeding any analysis of the drivers acting across different scales. An alternative solution is represented by meta-models and meta-analysis, which offer the possibility of reducing model output complexity by establishing a simplified mathematical relationship between the input and output of the system (Simpson et al., 2001). Their main aim is to replace complex numerical models by more parsimonious representations that provide a better understanding and faster analysis tools for optimisation and exploration, specifically when performing uncertainty or sensibility analysis (Simpson et al., 2001; Jia and Taflanidis, 2013; Saint-Geours, 2012; Ratto et al., 2012). Where possible, an elegant way to build meta-models is the approximation through an analytical model, which is fitted to the large-scale output and allows for simplification (Grimm and Railsback, 2005). Analytical solutions can provide insight from different aggregation levels, but their construction and use are not always unequivocal (see Johst et al., 2013). Spatial statistic techniques are potential candidates of great interest and should be further explored (Fritsch et al., 2020). For example, Jia and Taflanidis (2013) present a systematic implementation and optimisation of kriging meta-models for hurricane wave and surge prediction maps based on high-dimensional outputs to reduce complexity while preserving spatial dimension. In functional Magnetic Resonance Imaging analysis, Kang et al. (2014) show a meta-analysis approach to synthesise brain mapping information from images. Given brain activation maps, they propose a spatial point process approach to model peak activation locations, which were identified as local maxima of brain activation area, explaining the brain task involved.

Here, we show how spatio-temporally explicit outputs of population dynamics models in landscape ecology can be analysed through a meta-modelling approach. Such outputs are simplified to point patterns composed of individual positions, key events or significant hotspots defining local dynamics. The resulting patterns can be modelled as spatio-temporal point processes (STPP), and the pattern itself, or rather its structure, is the response variable that one seeks to explain through the structure of the spatial support, and its temporal changes, described through appropriately defined predictor variables (Diggle, 2003; Illian et al., 2012; Renshaw, 2015; Illian and Burslem, 2017). Point processes can be defined over continuous space and time, such that there is no need to work with fixed spatial and temporal units; they can be used for descriptive analyses and stochastic modelling of patterns. For example, Law et al. (2009) apply STPP tools by computing first- and second-order statistics, *i*.*e*., expected numbers of points, and of point pairs with given point-to-point distance, for characterising observed plant patterns; Gabriel et al. (2017); Opitz et al. (2020); Pimont et al. (2020) develop models for wildfire occurrences through STPPs to overcome challenges given by the multi-scale structure of data and by strong non-stationarities in space and time driven by weather, land-cover and land-use.

The main novelty of our work resides in the characterisation of spatio-temporal population dynamics through STPPs. As a case study application, we focus on the relationships among agricultural landscape structure and the dynamics of a pest and its natural enemy. A hierarchical framework is developed (Figure 1): (i) a stochastic landscape model, characterised by parameters determining the landscape configuration and composition, is constructed and simulated; (ii) a spatially explicit population dynamics model, characterised by parameters determining the pest-predator structure and its spatial heterogeneity, is constructed and simulated. We propose to represent spatio-temporally explicit outputs returned by this modelling chain as point patterns identifying space-time-indexed key events of pest dynamics, that we subsequently model by constructing and estimating statistical regression equations for multi-type STPPs. The response variables we aim to model are the occurrences and the magnitude of the pest density peaks. Response variables are explained by taking into account both global and local landscape features, species life-history traits, and the occurrences of pest inoculation, pest peaks and treatments in appropriately chosen spatio-temporal neighborhoods around the location and time where the response variable was observed. This approach allows us to investigate the role of landscape structure in influencing the point process intensity summarising the pest-predator dynamics, and we address two general questions: (1) How can landscape effects and population dynamics traits at different spatio-temporal scales be coupled? (2) What are the spatio-temporal relationships between pest inoculations, pest density peaks and landscape heterogeneity?

**Figure 1:**
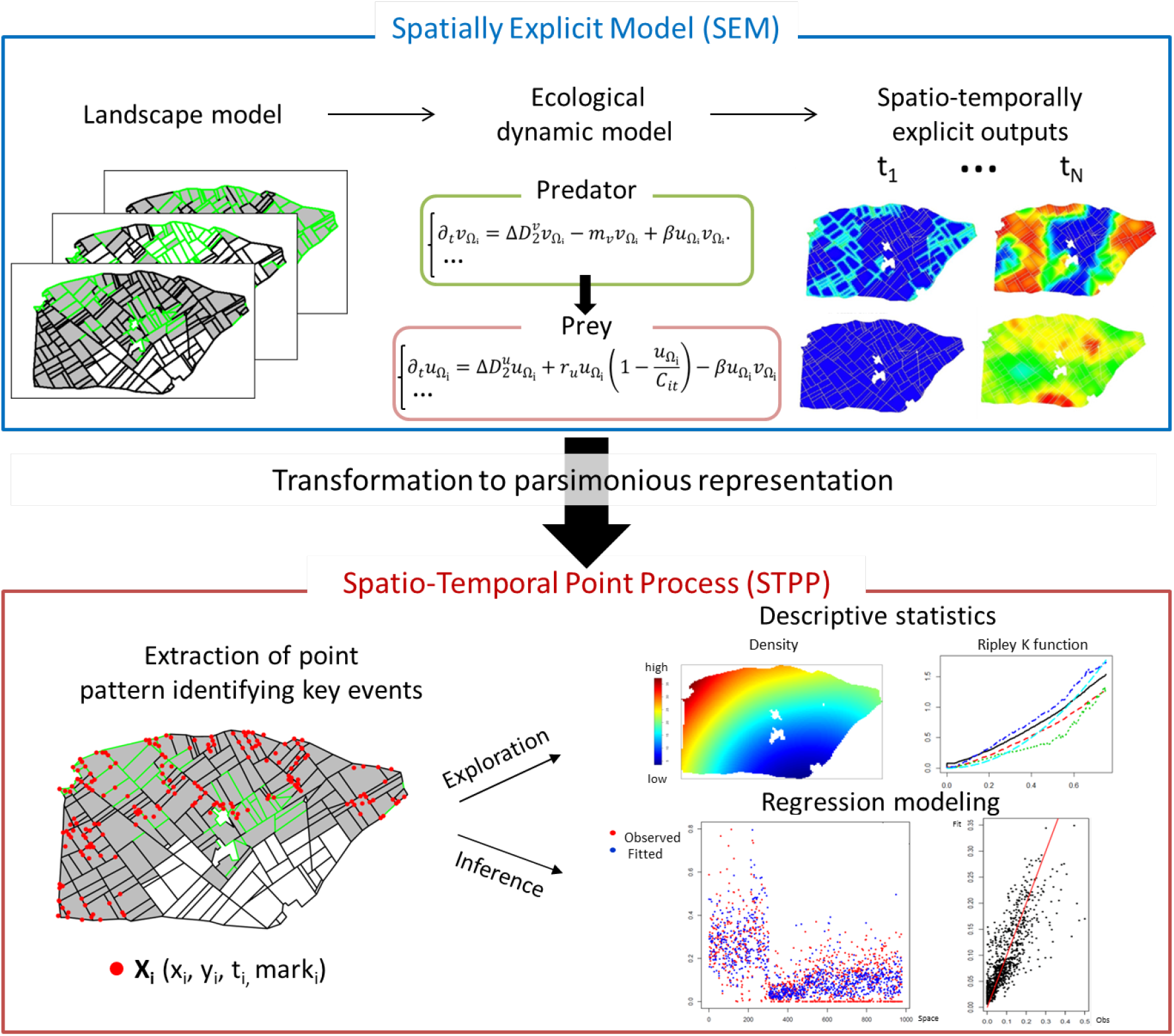
Overview of meta-modeling workflow.

## 2 Simulation models for landscape-pest-predator dynamics

### 2.1 Pest-predator models within agricultural landscapes

We model agricultural landscapes composed by crops, semi-natural areas and hedges through a stochastic landscape generator. Landscape simulations are the spatial support for a spatially explicit population model of auxiliaries and pests with opportune chemical treatments on pests. To couple the landscape complex and the spatially explict population model, we allow for dispersal both on agricultural fields and on hedge network (Figure 1). The agricultural landscape is composed of patches (*i*.*e*., polygons) and linear elements (*i*.*e*., segments) (Zamberletti et al., 2021). We generate a wide variety of structurally different composition and configuration scenarios for the allocation of crop over patches and of hedges over linear elements by varying representative parameters (*i*.*e*., crop and hedge proportion and their aggregation); details are provided in the Supplement. Within these generated spatial supports, we then simulate the dynamic of the codling moth (*Cydia pomonella*) pest and of one of its main predators, the family of ground beetles (*Carabidae*), in apple orchards. The pest-predator model is defined by a spatially explicit and density-based model of reaction-diffusion type (Roques and Bonnefon, 2016).

Codling moths respond strongly to the spatial distribution of orchards over landscapes (Tischendorf, 2001; Ricci et al., 2009). Franck et al. (2011) have found low genetic differentiation among codling moth populations over large distances, but mild genetic differentiation among populations collected on different host plants. In addition, insecticide treatments have strong effects on genetic differentiation resulting from spatial and temporal population size variations (Franck et al., 2011). This indicates that codling moths can disperse over large distances in agricultural landscapes, which supports the conjecture that hedges do not substantially impact their dispersal, such that insecticide treatments to break the pest dynamics are important. Thus, in the model, we assume that the pest can be encountered only in fields and that it has positive growth only in fields allocated with crop. In addition, field boundaries do not affect the pest population dynamics; *i*.*e*., the life cycle of *Cydia pomonella* is mostly based in apple orchards, and it perceives the landscape as a heterogeneous 2D environment. Finally, we impose the application of local insecticide treatments when the pest density exceeds a fixed threshold on average in a crop patch.

The presence of semi-natural areas, such as hedges, promotes the presence of pest auxiliaries (Maalouly et al., 2013; Thies and Tscharntke, 1999) by offering shelter and by providing complementary resources when pests are not present in fields (Lefebvre et al., 2017). Lefebvre et al. (2017) present a field study investigating the routine movement of arthropods among apple orchards and adjacent hedgerows. They found that there are frequent movements for foraging (to orchards) and for escaping treatments (to hedges), demonstrating the important influence of hedgerows on the presence of numerous predators in apple orchards. Thus, we consider that hedges form the main habitat of the predator. The predator can spill over from hedges to fields and there feed on pest in fields as an alternative resource. However, it is generally attracted to hedges, which are its preferred habitat, so that migration from fields to hedges is relatively high. The predator is known to be averse to moving outside its natural habitat; therefore, migration from hedges to fields is always lower than migration from fields to hedges (Lefebvre et al., 2017).

Details about the pest-predator dynamics among 1D and 2D elements are fully presented in Roques and Bonnefon (2016). All the parameters are shown in the Supplement. To fix parameter ranges, we had performed a sensitivity analysis in a preliminary step since observation data of pests and predators are not available (Zamberletti et al., 2021). Initially, the predator is present in all hedges at carrying capacity. The pest is introduced randomly in space and time. The time unit can be considered as the day. Overall, 172, 500 simulations were run by varying landscape and population parameter configurations (see parameter ranges in Table 1 of the Supplement), with 15 simulations for each configuration where parameters are fixed but landscape realisations are stochastic.

**Table 1:**
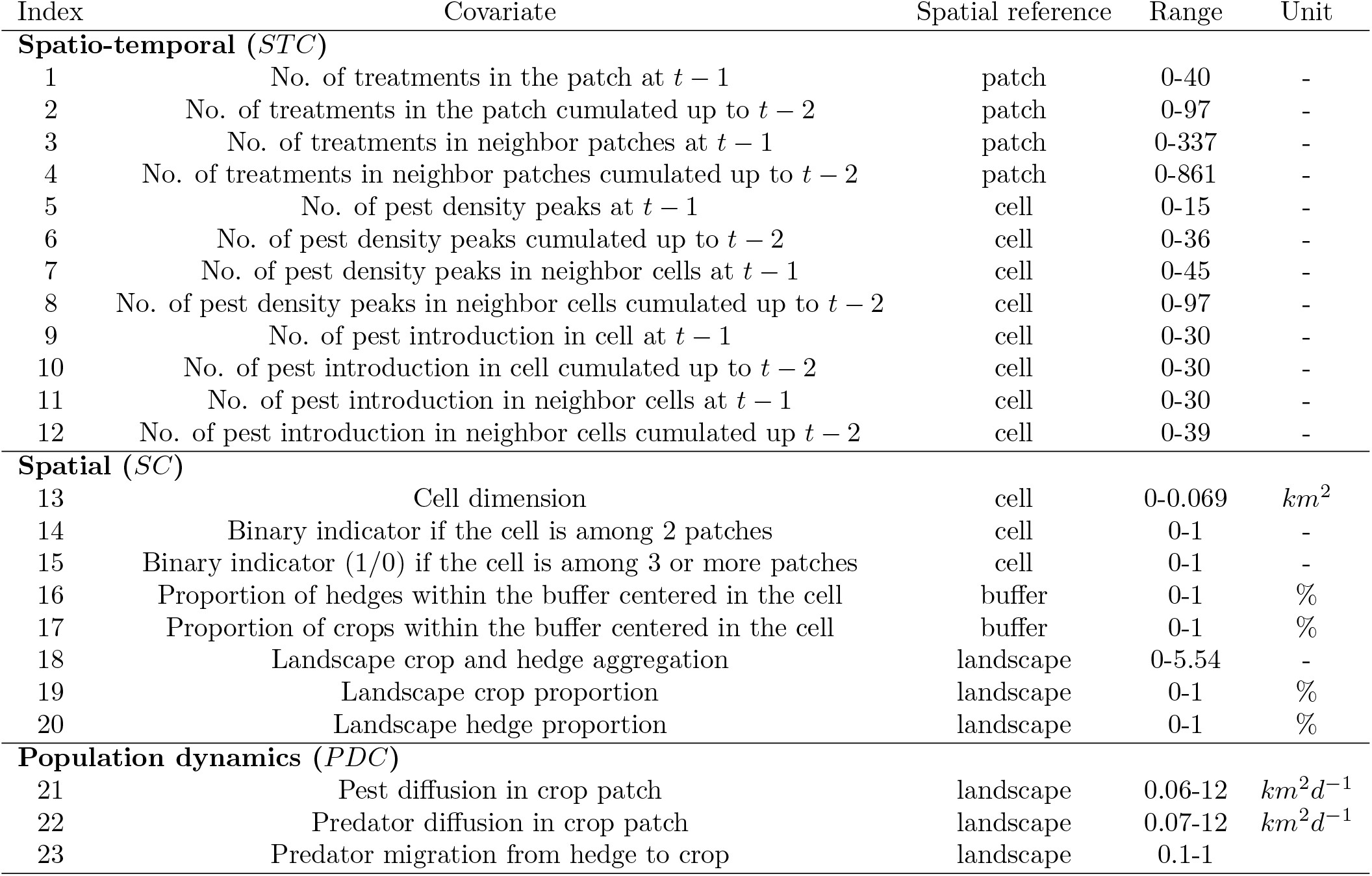
Covariates used in the space-time regression model of pest density peak patterns. The temporal unit *d* stands for *day*.

| Index | Covariate | Spatial reference | Range | Unit |
| --- | --- | --- | --- | --- |
| **Spatio-temporal (*STC*)** |  |  |  |  |
| 1 | No. of treatments in the patch at $t - 1$ | patch | 0-40 | - |
| 2 | No. of treatments in the patch cumulated up to $t - 2$ | patch | 0-97 | - |
| 3 | No. of treatments in neighbor patches at $t - 1$ | patch | 0-337 | - |
| 4 | No. of treatments in neighbor patches cumulated up to $t - 2$ | patch | 0-861 | - |
| 5 | No. of pest density peaks at $t - 1$ | cell | 0-15 | - |
| 6 | No. of pest density peaks cumulated up to $t - 2$ | cell | 0-36 | - |
| 7 | No. of pest density peaks in neighbor cells at $t - 1$ | cell | 0-45 | - |
| 8 | No. of pest density peaks in neighbor cells cumulated up to $t - 2$ | cell | 0-97 | - |
| 9 | No. of pest introduction in cell at $t - 1$ | cell | 0-30 | - |
| 10 | No. of pest introduction in cell cumulated up to $t - 2$ | cell | 0-30 | - |
| 11 | No. of pest introduction in neighbor cells at $t - 1$ | cell | 0-30 | - |
| 12 | No. of pest introduction in neighbor cells cumulated up to $t - 2$ | cell | 0-39 | - |
| **Spatial (*SC*)** |  |  |  |  |
| 13 | Cell dimension | cell | 0-0.069 | $km^2$ |
| 14 | Binary indicator if the cell is among 2 patches | cell | 0-1 | - |
| 15 | Binary indicator (1/0) if the cell is among 3 or more patches | cell | 0-1 | - |
| 16 | Proportion of hedges within the buffer centered in the cell | buffer | 0-1 | % |
| 17 | Proportion of crops within the buffer centered in the cell | buffer | 0-1 | % |
| 18 | Landscape crop and hedge aggregation | landscape | 0-5.54 | - |
| 19 | Landscape crop proportion | landscape | 0-1 | % |
| 20 | Landscape hedge proportion | landscape | 0-1 | % |
| **Population dynamics (*PDC*)** |  |  |  |  |
| 21 | Pest diffusion in crop patch | landscape | 0.06-12 | $km^2d^{-1}$ |
| 22 | Predator diffusion in crop patch | landscape | 0.07-12 | $km^2d^{-1}$ |
| 23 | Predator migration from hedge to crop | landscape | 0.1-1 |  |

### 2.2 Pest-predator spatio-temporal patterns

Simulations provide the spatio-temporal pest and predator densities. We characterise the influence of landscape spatio-temporal structure on the prey-predator dynamics by using point patterns. Following our modelling framework, we identify as events (i) the spatio-temporal treatment occurrence (*i*.*e*., pest threshold exceedance or pest peak) and (ii) the spatio-temporal pest introductions. For example, when pest threshold exceedance occurs in a patch, we apply a treatment in this patch and, to define the event episode as a point, we extract the time *t* of threshold exceedance, the pest density maximum in the patch with its Euclidean coordinates (*x, y*), and the average pest density over the patch. In Figure 2, two simulations are shown for different time steps, where the spatio-temporal occurrences of pest inoculations and treatments within different landscape allocations are highlighted. This example also illustrates the conjecture that the spatial hedge structure plays a role for pest dynamic by influencing its evolution jointly in space and time. Deeper exploratory quantitative analyses of spatio-temporal relationships between different types of points are proposed in the Supporting information, while we focus on statistical model-based analyses in what follows.

**Figure 2:**
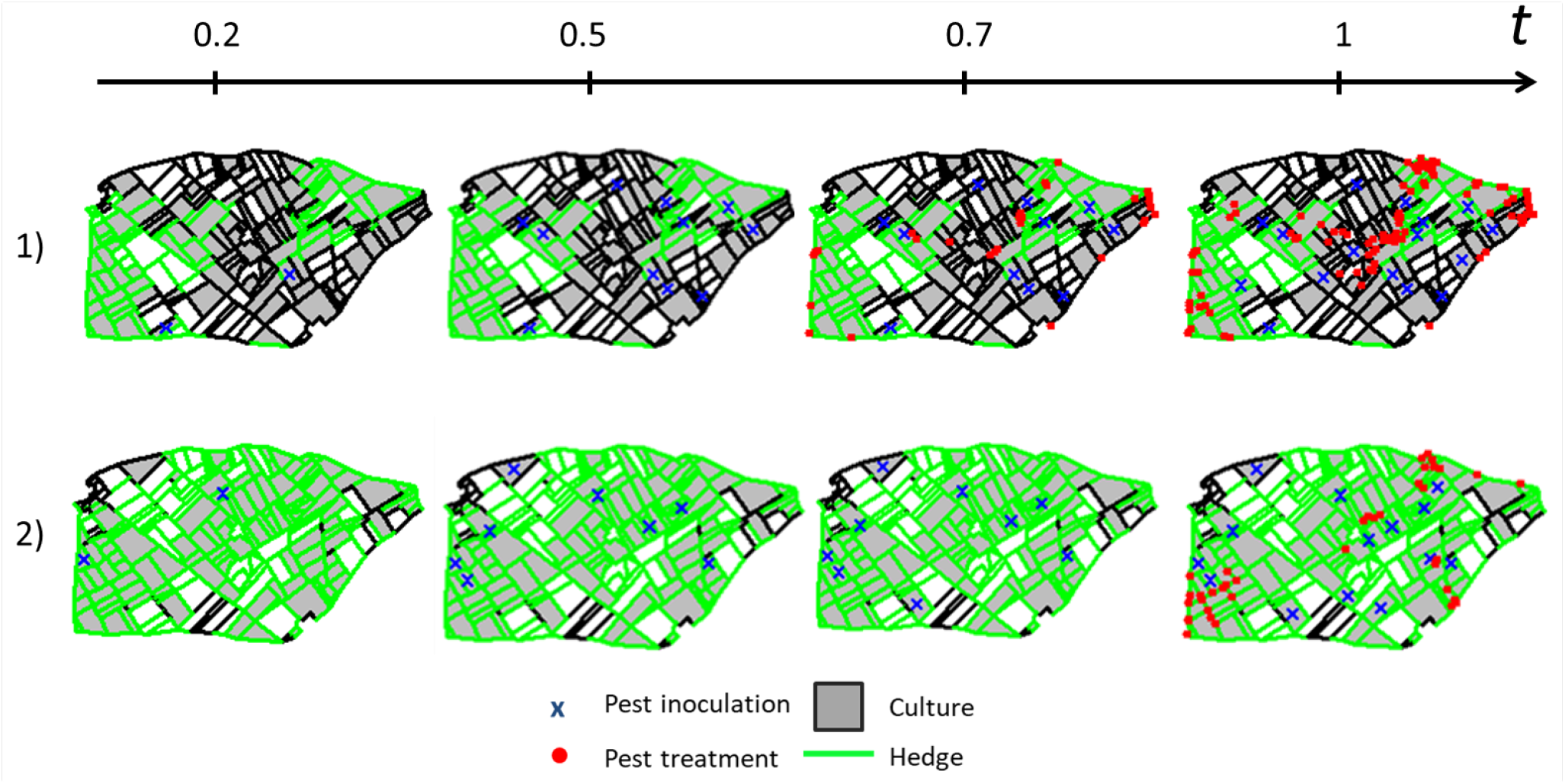
Two simulation examples (by row) illustrating the spatio-temporal pest dynamics depending on landscape structure through pest inoculations, and through pest density peaks after threshold exceedances.

## 3. Methods: STPP-based analysis of pest-predator dynamics

### 3.1 Pest density as STPP

Point patterns representing individual or event distributions in space and time can be modelled as STPPs (see Diggle (2003); Illian et al. (2008); Baddeley et al. (2015) for formal definitions). Each point can be endowed with additional qualitative or quantitative information defined as a “point mark”. In our application, the pattern of events is defined by the coordinates in space and time of pest peaks with both qualitative (pest inoculation) and quantitative marks (pest maximum density). Thanks to the theory of STPPs it is possible to analyse the point distribution properties locally in space and time, and to estimate models for predictive purposes (*e*.*g*., number of events, point-to-point correlations, and distribution of their numerical or categorical marks). We focus on modelling the point process intensity function (local point density) (Illian et al., 2013). Our modelling goal is to predict the intensity of pest density peaks and the associated values of maximum pest density, and explain their variability in space, through time and across different simulations. We divided the spatial domain in a relatively large number of small cells, and we assume a homogeneous point process intensity within each cell during each interval of time. The spatial discretisation we use is shown in Figure 3, and background on its structure and construction is provided in the Supplement.

**Figure 3:**
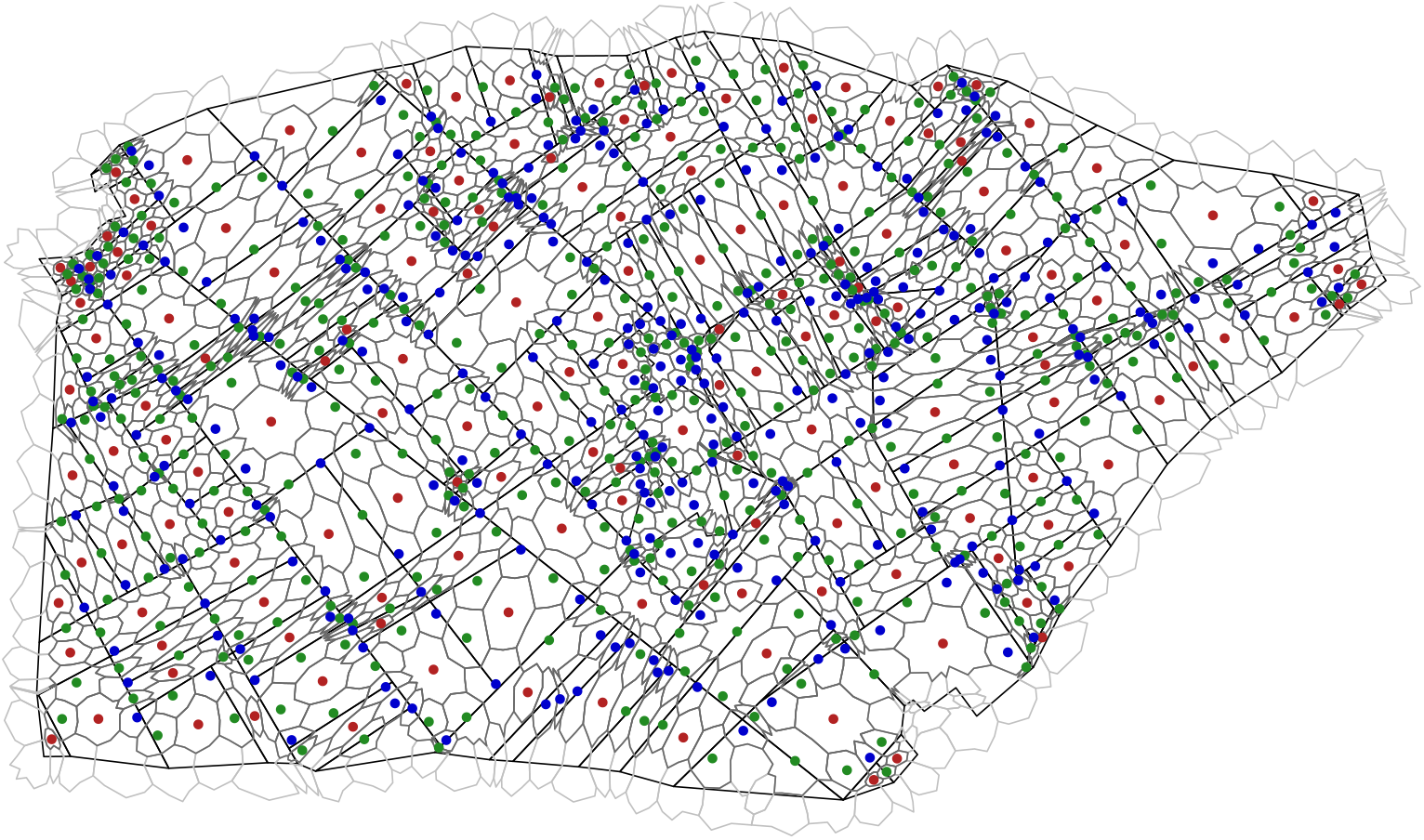
Spatial discretisation of the regression models. Complete mesh discretisation (light grey), mesh cells used in the analysis (dark grey), landscape patches (black). Cell centroids of different colour refer to different cell types: cell in patch center (red), cell connecting exactly two patches (green), cell connecting more than two patches (blue).

### 3.2 Pest density peak meta-modelling

For predicting the intensity of pest density peaks and associated values of maximum pest density, we develop and estimate regression equations for multi-type STPPs. Both global and local landscape features, species life-history traits, and the occurrences of pest introductions, pest peaks and treatments are used as covariate information. We construct two separate generalized linear model (GLM) formulas as meta-models that incorporate the available covariate information. Response variables and covariates are evaluated over each spatial cell (Figure 3) and time step. The spatio-temporal (*STC*), spatial (*SC*) and population dynamics (*PDC*) covariates put the spatio-temporal event patterns, landscape structure and population dynamics into relation:

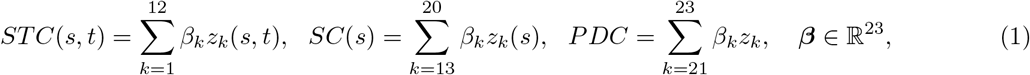

The ***β*** vector gathers the covariate coefficients to be estimated separately for each model, and the values *z*_*k*_ are covariates summarised in Table 1 and provided for each space-time cell. More information on their selection and computation is given in the Supporting information, as well as residual analysis to evaluate the predicted values obtained by the GLMs.

#### 3.2.1 Meta-model for the occurrence intensity of pest density peaks

To model the occurrence intensity of pest density of pest peak points, we consider a GLM with Poisson response, *i*.*e*., we combine a log-link function with a Poisson response distribution:

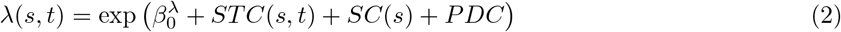

with global intercept 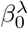 and coefficients of the other variables to be estimated. The value *λ*(*s, t*) represents the average number of pest peaks occurring in a unit of space and time around the point (*s, t*), and is assumed to be constant within each cell of the mesh during each time interval of 0.1.

#### 3.2.2 Meta-model for magnitudes of pest density peaks

To model the maximum pest density value associated with each pest peak point, we consider a log-Gaussian GLM, *i*.*e*., we combine a log-link function with a Gaussian response distribution:

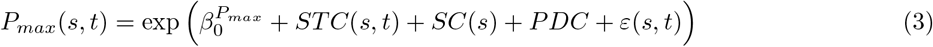

with global intercept 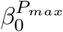 and coefficients of the other variables to be estimated, where *P*_*max*_(*s, t*) is the maximum pest density value associated to the point where the treatment is applied conditional to the occurrence of such a point. The term *ε*(*s, t*) ∼ 𝒩 (0, *σ*^2^) corresponds to the spatially and temporally independent and identically distributed Gaussian error terms.

## 4 Results: spatiotemporal drivers of pest hotspots in pest-predator agroecological system

We present main results obtained by estimating the GLMs in Equations 2 and 3. Additional results of a covariate correlation analysis and of residual analysis are reported in the Supporting information; they show that the models defined in Equations 2 and 3 appropriately capture the spatio-temporal variability of the observed data (*i*.*e*., population dynamic model outputs).

The estimated GLM coefficients for the models in Equations 2 and 3 are summarized in Figure 4. Prior to estimation, covariates have been normalised to empirical mean 0 and variance 1 to compare more easily the magnitudes of estimated effects.

**Figure 4:**
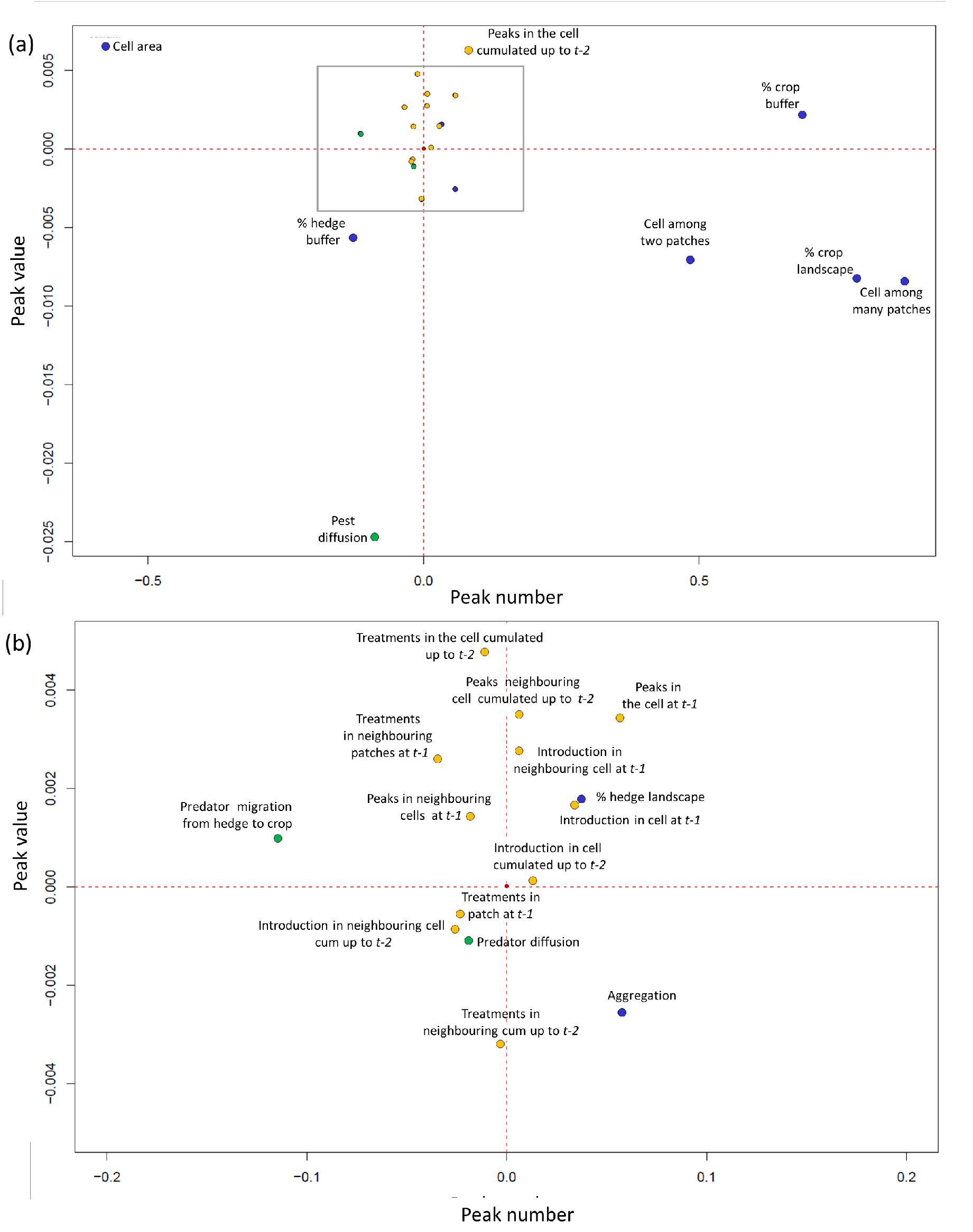
Estimated regression coefficients for the models of peak occurrence intensity (*x*-axis) and the model of the peak value (*y*-axis). Dot colours indicate covariate types: *STC* (orange), *SC* (blue), *PDC* (green).

We first discuss the strongest effects corresponding to points outside the inner rectangle in Figure 4a. The strongest positive effects on the number of pest peaks arise for covariates favouring pest dynamics.

Specifically, crop coverage at local scale (*i*.*e*., in the buffer) and at global scale (*i*.*e*., in the whole landscape) favours the abundance of suitable habitat for pests, which can easily spread and find resources. Regarding the pest peak value, the cell size has the strongest positive contribution. An explanation is that the pest density is likely to be highest where the inoculation takes place, and a large cell is more often inoculated than a smaller cell. By contrast, cell dimension contributes the strongest negative effect on the number of peaks, since peaks tend to concentrate in the periphery of the patches, thus in cells containing borders among different patches.

Pest diffusion has the strongest negative effect on pest peak values, it may be due to a dilution effect. In addition, since high pest diffusion allows the pest to easily move, pest population tends to spread homogeneously over the whole landscape. Therefore, few local hotspots arise, and the pesticide threshold is less often exceeded. Both response variables related to pest peaks are also strongly reduced by local predator presence, which in turn is mainly driven by a high local presence of hedges. The spatio-temporal covariate group (STC) shows generally weaker effects on pest dynamics, except for the local cumulated number of pest peaks during earlier time intervals. It positively influences the number and the value of pest peaks since pests are already present at high density in the surrounding area if there have been peaks during earlier intervals. Such locations may have characteristics that make them particularly pest-prone and favourable for pest dynamics.

The zoom in Figure 4b shows covariate effects with a lower magnitude. High numbers of pest peaks along with high peak concentration values (top-right quadrant in Figure 4) are relatively strongly favoured by the presence of previous peaks in the same cell or in the surrounding ones (both at *t* − 1, and cumulated up to *t* − 2). Similarly, an elevated number of introductions in neighbouring cells leads to high pest concentration due to pest spillover. On the other hand, the application of treatments locally in the patch or in neighbouring patches at previous time steps leads in general to a decrease of both the number and the concentration value of peaks.

Results show a negative effect of hedge proportion in the buffer on pest activity. However, there also arises a weaker but positive effect of the hedge proportion over the whole landscape, which may appear counter-intuitive at first glance. Since response variables are evaluated at cell scale, having a large hedge proportion in the whole landscape but a low proportion of hedges in the buffer clearly results in a concentration of pest where hedges are missing. In addition, hedges help to keep the pest below the treatment threshold and therefore favour its propagation through the landscape (see Zamberletti et al. (2021)); therefore, the pest may reach areas of lower predation pressure more easily and pull out. In addition, our model shows that the landscape aggregation has a weak positive effect on peak occurrence numbers at cell level. Pest density threshold exceedances occur homogeneously over large areas of contiguous crop, but these peaks are of relatively small magnitude because hotspots with high pest clusters and concentration do not build up. Predator spillover (*i*.*e*., movement from hedge to field) results in a decrease of the number of threshold exceedances, but it may increase pest peak values since the predators are not homogeneously present in the patches and over the whole landscape. Predators have stronger influence near hedges (*e*.*g*., in cells overlapping different patches) but less in the center of the patch (central cells).

## 5 Discussion

In this work we propose post-model scaling using regression meta-models based on marked STPPs. This approach enabled us to assess and compare the contribution of different spatio-temporal covariates and life-history traits to the direction and strength of variation in crucial events of population dynamics issued from spatially explicit models. The use of statistical regression meta-models makes our approach flexible and easy to implement, while numerous and diverse covariates describing local and global characteristics can be incorporated. We applied our methodology to the outputs of a SEM describing the biological control in agricultural landscapes of a crop pest by its natural predator. We found significantly different effects of landscape structures at various spatial scales on the population dynamics patterns.

The adaptation of our approach of defining a marked STPP meta-model may be relevant and insightful in various contexts. Examples are occurrence locations and times of earthquake epicentres (Lombardo et al., 2019), wildfires (Opitz et al., 2020), epidemiological outbreaks (White et al., 2018a), biodiversity hotspots and species distribution (Soriano-Redondo et al., 2019), pollutant concentrations (Lindström et al., 2014) or local maxima or minima in meteorological events (Heaton et al., 2011). In most ecological process space and time are closely intertwined and not separable as in our case, where pest introductions and subsequent peaks depend on local temporal dynamics driven by local spatial structure. Thus, here, we designed our approach to allow for joint analysis of spatial and temporal scales. For ecological processes related to those we study, White et al. (2018a) addressed how landscape structure impacts simulated disease dynamics in an individual-based susceptible–infected–recovered model. They quantified disease dynamics by outbreak maximum prevalence and duration, coupled with landscape heterogeneity defined by patchiness and proportion of available habitat. They find that fragmentation promotes pathogen persistence, except for simulation with high conspecific density, slower recovery rates and larger perceptual ranges, where more complex disease dynamics emerged; the most fragmented landscapes were not necessarily the most conducive to outbreaks or pathogen persistence. Our work has similar thrust by exploring the effect of landscape heterogeneity on pest density peaks. However, by taking advantage of the STPP modelling, we focus on spatio-temporal positions of peaks, and we investigate which factors locally influence occurrence intensity and magnitude of these events. The meta-model allowed us to depict complex spatial dynamics and patterns even if multiple processes occur at competing scales (White et al., 2018b). To assess fine-scale biodiversity, Azaele et al. (2015) captured species patterns through correlations among different species’ abundances across sample plots. Therefore, they used counts over spatial units (*i*.*e*., plots), determined by the sampling design and leading to relatively large counts, and they contrasted their results with common species–area curves (Fritsch et al., 2020). They concluded that this mathematical framework provides a common language to link different spatial scales. Our approach goes beyond a purely descriptive “geostatistical” analysis since we take into account the space-time position of each of the points as well as their relationships with nearby key elements. This representation parsimoniously summarises spatially continuous dynamics into discrete occurrences of spatio-temporal key events and allows modeling them for explanatory and predictive purposes. Our regression model for occurrence intensities also aggregates individual events, but we work with relatively small counts by choosing appropriate, problem-specific space-time units.

Ecosystem patterns and processes can cover a wide range of space and time, and they depend on multiple drivers acting over different scales (Fritsch et al., 2020). Problematic loss and the lack of information may arise in procedures of scaling-up or scaling-down when coupled with the complexity of the involved systems. Our work strikes a pragmatic balance with respect to the inevitable trade-off between model simplicity, to obtain clear insights into important factors, and model complexity, to achieve a more complete and realistic representation of the system (Lacy et al., 2013). Spatio-temporal meta-models present a flexible solution by capturing the functional linkages between model components. They show potential to reveal properties in ecological systems that are difficult to identify when considering only the complex model output with large data volumes as a whole (Lacy et al., 2013). Our STPP model allowed for a relatively complex spatio-temporal local analysis of system dynamics. It therefore provides insights into the role of different effects and takes process-specific scales into account by using categorical or numerical marks. Through statistical inferences it becomes possible to identify significant relationships of key events with their drivers focusing on biotic interactions, habitat heterogeneity and spatio-temporal stochastic effects predictions (Baddeley et al., 2015).

A large body of literature on meta-models (or surrogate models, or emulators) in various disciplines focuses on Gaussian processes or machine-learning techniques (*e*.*g*., Forrester et al., 2008; Kleijnen, 2015), whereas our work highlights the potential of point-process-based approaches for dynamical systems. This novel way of conducting meta-analyses is applicable to various collections of relevant events arising in dynamical processes available at high spatio-temporal resolution. We emphasise that our methods leverage spatio-temporal and multivariate point pattern techniques, while the state-of-the-art in point pattern analyses deals mostly with purely spatial patterns or does not well represent the temporal dimension (Wiegand et al., 2017). Our extensions are well-suited for spatio-temporal mechanisms and population dynamic parameters where the assessment of their relative and joint role is crucial for characterising emerging diversity patterns.

We have constructed a collection of predictor variables in which spatio-temporal covariates (STC) contribute spatio-temporally structured information, such as the number or magnitudes of previous or concomitant events around a given location and time, to convey information related to the local evolution of pest dynamics. In a similar context, Le Gal et al. (2020) highlighted the important influence of the interplay between the landscape structure and the timing of CBC measures on the delivery of pest control services. They showed that increased semi-natural habitat proportion at the landscape level enhances the visitation rate of pest-colonised crop cells, but it also reduces the delay between pest colonisation and predator arrival in the crop fields. In our model, we have opted for simulating the time and position of pest arrival according to a Poisson process with intensity proportional to crop area. We found that locations showing frequent and high density peaks in previous time steps are likely to incur new peaks. On the other hand, local previous treatments in a patch negatively influence the dynamics since they efficiently reduce the pest density in this patch. Introductions of pest act as an accelerator of local pest dynamics, and after a short period we often assist to both high frequency and high magnitudes of peaks in the surrounding fields.

Spatial covariates (SC) in our regression meta-models are time-invariant landscape characteristics that may influence pest peaks. Crop proportion is the main driver for pest in our models, and leads to a clear positive response of pest insects to increasing cover of a suitable crop (Ricci et al., 2019; Rand et al., 2014; Zhao et al., 2015; Avelino et al., 2012; Tscharntke et al., 2007). Our results show that considering it at local scale or at global scale leads to different peak patterns. When crop aggregation and percentage coverage are high in the whole landscape, exceedance events of pest density are relatively homogeneously spread over the area with generally relatively low pest density values throughout. Instead, when high crop coverage is only local (*i*.*e*., in the buffer), the resulting pattern shows a locally higher number of exceedance events with high peaks; pests find their preferred habitat in a more limited space and tend to concentrate there. Zamberletti et al. (2021) showed that in landscapes with strong aggregation of crop fields the area of contiguous crop may cause a dilution effect, with a positive effect on pest population, a negative effect on treatment occurrence, and a positive effect on the treatment numbers in the whole landscape. Therefore, if treatments are necessary in a patch, they tend to arise in relatively high numbers over the full observation period. Hedge distribution and proportion can be viewed as a proxy for predator presence and reveal when predators may play a role in reducing pest density (Bianchi et al., 2006; Tscharntke et al., 2007). The effects attributed to semi-natural habitat (*e*.*g*., hedges) are ambiguous with both positive, negative or neutral impacts on CBC (Chaplin-Kramer et al., 2011; Karp et al., 2018). In our models, total hedge proportion has a small but positive effect on both the number and the magnitude of peaks. A reason could be that the global proportion of hedges does not inform about hedge connectivity and distribution (*e*.*g*., homogeneously or in clusters). If there is a high hedge coverage, predators are expected to be homogeneously distributed in the landscape, thus stabilising the pest population and potentially reaching an equilibrium in the whole landscape for pest and predator density. However, this does not imply that pest density remains under the treatment threshold; it could happen that other parameters influence its dynamics by favouring pest population (*e*.*g*., crop coverage or pest growth rate) or decreasing predator presence in field (*e*.*g*., mortality, spillover from hedge). This results in a homogeneous predator presence that is not sufficient to prevent pest density from exceeding the threshold. In our model, another reason could stem from statistical confusion in the regression models between the effects of global hedge proportion and global crop proportion since the simulated landscape model tends to position hedges more often in crop areas than in the rest of the landscape. However, when focusing on local buffers around a cell, local hedge structure, and the resulting predator concentration, play a bigger role by reducing both number of pest peaks and their magnitude.

Population dynamics covariates (PDC) in our models are related to species traits. Here we consider the effect of varying population parameters related to species mobility in the environment. We focus on how the structure of landscape elements influences species spread with respect to the studied events. We find that predator diffusion ability over the landscape is fundamental to reduce the presence of pest. Interestingly, we do not notice the same effect for predator migration speed from hedge to field. This predator trait acts strongly at locations close to hedges, *i*.*e*., around patch borders, with a strong decrease in the number of peaks, while the peak value is not affected but is high mainly in the patch core areas.

In the agro-ecological context, our analysis aids prediction and management decisions. For example, improved understanding of local spatio-temporal relationships and dynamics helps to schedule specific local control strategies by targeting the locations that frequently suffer from pest peaks and the moments when local control strategies can be expected to be most efficient to control pest dynamics.

## Notes

### Competing Interest Statement

The authors have declared no competing interest.

