## Supplementary material for "Spatio-temporal point processes as meta-models for population dynamics in heterogeneous landscapes"

#### 1 Exploratory spatio-temporal analysis

We give an example of the exploratory analysis of relevant behaviour and effects when we are close in both space and time to some reference or trigger event (*e.g.*, when studying what happens after an inoculation, or after a pest peak) and of relationships with the landscape structure. This analysis aims to highlight the importance of jointly considering the space and time dimensions in our analysis.

The spatio-temporal structure of pest peaks is explored by evaluating the pest peak occurrence intensity after a peak or after an inoculation. For each group of 15 repetitions of the 172,500 simulations, we select the first temporal pest peak or the first inoculation, and we then compute the spatial and temporal distances to all the other treatments later in time. Then, we use such distances as point in the $xy$ plane and visualise the resulting cloud of points intensity. We run analyses where we either pull together all the simulations, or we run analysis on subsets of data by dividing them depending on key parameters. In Figure 1, we report the logarithm of the occurrence intensity of treatments with respect to the spatio-temporal distances among $1^{st}$*peak-peaks* ( Figure 1a) and $1^{st}$*inoculation-peaks* (Figure 1b), where we consider the following data subsets: all the simulations together (first column); by splitting the simulations depending on the *% of crop* in the whole landscape (low and high in second and third columns, respectively); by splitting the simulations depending on *pest diffusion in crop* (low and high in forth and fifth columns, respectively).

The plots for $1^{st}$*peak-peaks* allow us to assess how rapidly the pest dynamic is able to recover after the first peak and the associated application of a treatment within a patch to exceed again the treatment threshold at a later stage, which also involves the re-colonisation from the neighbouring patches (Figure 1a). When considering all the simulations, given a temporal distance, the peak intensity decreases moderately with short spatial distance and more rapidly as the spatial distance increases. On the other hand, given a spatial distance, there is a increasing trend of pest peak intensity until reaching a maximum value at a certain temporal distance, and then it decreases. The temporal range with high values is wider for shorter distances and shrinks as the spatial distance increases. In general, we observe a maximum intensity around the time step $t = 0.4$ at all spatial distances. This behaviour can be explained by the fact that pests need time to disperse and reach the threshold density above which the treatment is applied; and, at a short distance the pest threshold would be exceeded more quickly than at a longer distance. Given a temporal distance, pest density will be higher for shorter spatial distances. This behaviour changes when looking only at simulations with small % *of crop*, since pests have not a lot of space in their preferred habitat. Thus, they group in a small spatial range of suitable habitat. Consequently, the maximum intensity is reached at relatively short spatial and temporal distances. By contrast, for high *pest diffusion in crop*, we observe a larger temporal range at which the maximum pest peak intensity is observed, which is due to the fact that pests disperse rapidly, thus are able to spread in shorter time than on average for all simulations.

The plots for the $1^{st}$*inoculation-peaks* analysis allow us to assess the capability of the pest population to develop and colonise the surrounding landscape (Figure 1b). For all the simulations, the maximum occurrence intensity of treatments depends mostly on the spatial distances with a decreasing trend at small to moderate spatial distances; therefore, the closer we are to the inoculation point, the more pest peaks above the treatment threshold will occur. More specifically for a small % *of crop*, the maximum occurrence intensity is always found for relatively small spatio-temporal distances, where for small temporal distances relatively high occurrence intensities persist over all spatial distances. This behaviour could be due to the fact that pests cannot disperse over larger surfaces and therefore may locally accumulate very fast. For high *pest diffusion in crop*, the trend is completely different; we note generally higher intensity values with respect to the other cases, and the maximum intensity is realised for short spatial distances for all temporal distances. However, when the spatial distance increases and for short temporal distance, we observe relatively small pest peak occurrence intensities, due to the fact that pests spread quickly and may thus easily reach other habitats, which lowers the local pest density around the inoculation point and keeps it under the treatment threshold. At longer temporal distances, the pest propagation causes the threshold to be exceeded quite frequently.

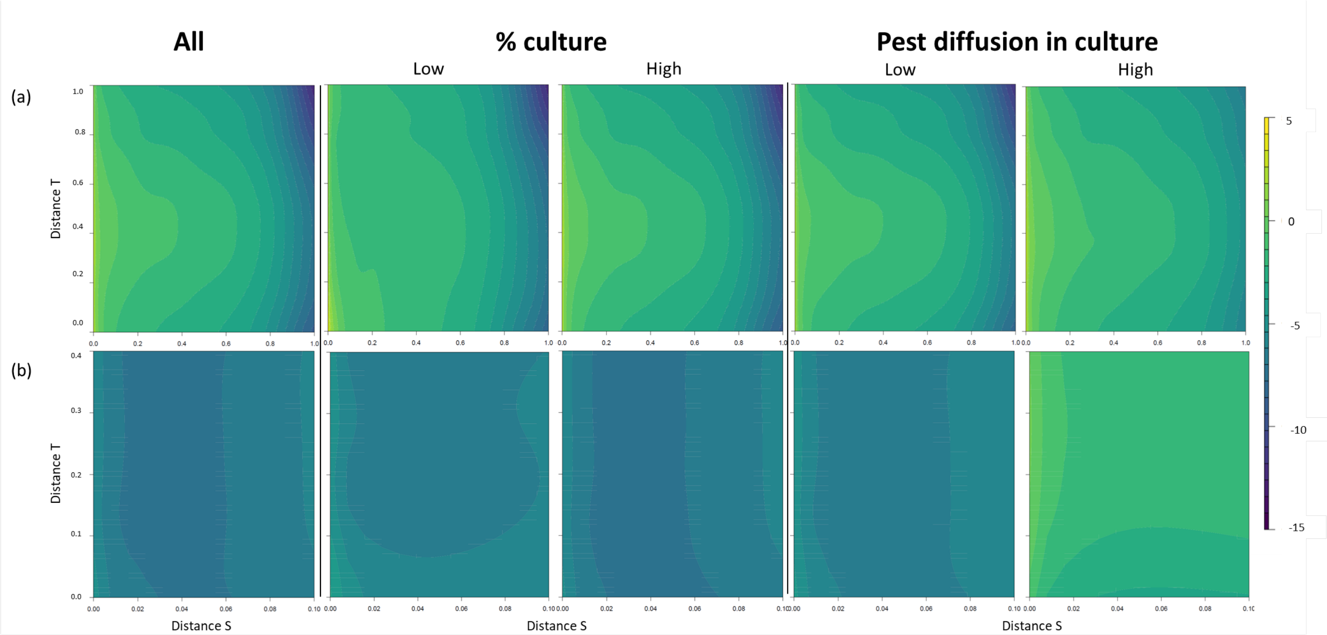


Figure 1: Spatio-temporal visualisation of pest peak occurrence intensity as a function of the spatial distance ($x$ axis) and the temporal distance ($y$ axis) with respect to the first pest peak in the simulation (Panel a) or with respect to the first inoculation in the simulation (Panel b); for all the simulations together (first column), or depending on low/high *% of crop* (second and third columns) and low/high *Pest diffusion in crop* (fourth and fifth columns). The colour bar displays the log-occurrence intensity.

### 2 Landscape model

We provide details about the landscape model, which was developed in Zamberletti et al. (2021). As fixed spatial support, we use an observed agricultural landscape and represent it through a vectorial approach by T-tessellation. It results in 188 polygons with a total of 577 edges. For simulating the spatial allocation of land-use categories in the T-tessellation, stationary Gaussian random fields (GRFs) are simulated in the landscape (with mean 0 and variance 1). We then fix a threshold on the values of the Gaussian field simulated at locations that represent the barycenter of a specific landscape element type (*e.g.*, polygons), and we allocate the landscape elements with one of two possible categories *e.g.*, crop, no crop) depending on the value of the GRF being below or above the threshold. The correlation parameters of the GRFs control the strength of spatial dependence, which, in turn, governs the clustering strength of landscape elements, such as surface elements and linear elements. The commonly used valid spatial correlation functions are restricted to nonnegative correlations, such that GRFs are useful to generate clustered or independent structures in space. It would be more difficult to obtain regular structures, *e.g.*, repulsive structures between neighboring elements. To simulate landscape configurations, GRFs are thresholded at a fixed quantile (*e.g.*, observed 70% quantile, where the probability level is one of the parameters of the model) to define presence (if above threshold) or absence (if below threshold) of crops or of hedges, respectively. For each type of elements of the landscape geometry (*i.e.*, , segments and polygons), a separate GRF is defined to assign an land-use type controlling its spatial aggregation and proportion (*i.e.*, $W_1(s)$ and $W_2(s)$ for polygons and segments, with exponential correlation function given as $corr(s_1, s_2) = \exp(-\text{dist}(s_1, s_2)/\phi_i)$, $i = 1, 2$, respectively). Next, correlation between allocation of crop to patches and of hedges to edges can be generated by inducing correlation between their respective GRFs, here using the idea of a linear models of co-regionalization. Initially independent GRFs for landscape geometries can be linearly combined through correlation parameters (Equation 1, 2 and 3) to obtain the two dependent GRFs used to determine the final landscape configuration coupling surface and linear element allocation:

$$W_c(s) = \rho_c W_1(s) + \sqrt{1 - \rho_c^2} W_2(s) \tag{1}$$

$$W_h(s) = \rho_h W_1(s) + \sqrt{1 - \rho_h^2} W_2(s) \tag{2}$$

For a simpler and more parsimonious formulation, we fix $\rho_h = 1$ such that $W_1$ defines the GRF used for hedges, and we allow for $\rho_c \in [-1, 1]$ to control the correlation between $W_h$ and $W_c$ (Zamberletti et al., 2021). Moreover, we use the same spatial correlation range $\phi_1$ in both of the initial fields. Then, the cross-correlation function between the GRFs for hedges and crops is as follows:

$$\text{Corr}(W_h(s_1), W_c(s_2)) = \rho_c \exp(-\text{dist}(s_1, s_2)/\phi_1) \tag{3}$$

Specifically, for correlation of hedges and crops at the same location x, we obtain $\text{Corr}(W_h(x), W_c(x)) = \rho_c$.

### 3 Population dynamics model

We report the population dynamics model to provide more details with respect to the narrative of the paper. The model is the same of Zamberletti et al. (2021). Population dynamics are described by a spatially explicit predator-pest model based on a system of partial differential equations. Our model is built on an earlier developed approach that considers both 2D diffusion on surface elements and 1D diffusion on linear elements (Roques and Bonnefon, 2016).

- Predator model structure:
  - 1D landscape elements:
    Linear 1D elements of the landscape matrix are denoted by $h_i$. A 1-dimensional reaction-diffusion model on linear elements is defined for the predator $v_{h_i}$:

$$\begin{cases} \partial_t v_{h_i} &= \partial_{xx} D_1^v v_{h_i} + r_v v_{h_i}(1 - \frac{v_{h_i}}{K_{h_i}}) \quad \text{if the edge } h_i \text{ carries a hedge,} \\ v_{h_i} &= 0 \quad \text{otherwise,} \end{cases} \tag{4}$$

    where $D_1^v$ is the diffusion parameter of the predator along the hedges, $r_v$ is the intrinsic growth rate of the predator, and $K_{h_i}$ is the carrying capacity of the hedge $i$.
  - 2D landscape elements:
    Polygon-shaped 2D fields are denoted by $\Omega_i$. The population density of predators $v_{\Omega_i}$ in each field is modelled by a reaction-diffusion equation with mobility parameter within field $D_2$, predation rate $\beta$, and mortality $m$:

$$\partial_t v_{\Omega_i} = \Delta D_2^v v_{\Omega_i} - m v_{\Omega_i} + \beta u_{\Omega_i} v_{\Omega_i}. \tag{5}$$

- Prey (i.e. pest) model structure:
  - 1D landscape elements:
    We make the assumption that edges do not host the pest $u_{h_i}$, and that they do not directly modify their population dynamics:

$$u_{h_i} = 0 \quad \text{for all } i. \tag{6}$$

– 2D landscape elements:
The pest $u_{\Omega_i}$ is assumed to live only in fields. In addition, the crop fields represent a source of pest, whereas the non-crop fields are a sink for the pest. In the absence of dispersal from fields hosting the crop, the pest population vanishes in fields hosting the non-crop type. A chemical treatment is applied to a given crop field when the pest population in that field reaches a given threshold. The bidimensional reaction-diffusion model is defined as follows:

$$\begin{cases} \partial_t u_{\Omega_i} &= \Delta D^v_{u_{\Omega_i}} u_{\Omega_i} + r_{u_{\Omega_i}}(1 - \frac{u_{\Omega_i}}{C_{it}}) - \beta u_{\Omega_i} v_{\Omega_i} \quad \text{for crop,} \\ \partial_t u_{\Omega_i} &= \Delta D^v_{u_{\Omega_i}} u_{\Omega_i} - m_{u_{\Omega_i}} - \beta u_{\Omega_i} v_{\Omega_i} \quad \text{for semi-natural,} \end{cases} \tag{7}$$

where the carrying capacity $C_{it}$ of the field $i$ can change in time due to chemical treatments: $C_{it} = K_{\Omega_i}$, if no chemical treatment is applied, and $C_{it} = \frac{200}{K_{\Omega_i}}$ for short period of time after the chemical treatment is applied.

The dynamics described by equations 4 to 7 are coupled to define predator-pest dynamics over landscapes. Moreover, the fluxes of individuals between 1D and 2D elements of the landscape are defined as follow:

- Edges (with or without a hedge) do not affect the pest population dynamics, *i.e.*, the pest perceives the landscape as a heterogeneous 2D environment without 1D effects of linear elements.
- Edges without a hedge do not affect the predator population dynamics, *i.e.*, two fields separated by an edge but without a hedge will be perceived as a unique (potentially heterogeneous) 2D element by the predator.
- The predator is attracted by hedges, thus migration from fields to hedges is relatively high.
- The predator could potentially have an aversion to move outside its natural habitat; therefore, migration from hedges to fields is always lower than migration from fields to hedges. Finally, we considered reflecting conditions on the boundaries of the landscape, meaning that in- and out-fluxes between the landscape and its surrounding environment are equal.

Dynamics among 1D and 2D elements are fully presented in (Roques and Bonnefon, 2016). The parameter that controls the predator movement between linear elements and fields is $\rho_{12}$. All the parameters of predator and pest dynamics are shown in Table 1.

Table 1: Parameters of the population dynamics model.

| **Parameters** | **Description** | **Range** | **Units** |
|---|---|---|---|
| **For landscape model** | | | |
| $\phi$ | Aggregation of hedges and crops | $[5.55 \times 10^{-2} - 5.55]$ | - |
| $P_c$ | Proportion of crops | $[0 - 1]$ | - |
| $P_h$ | Proportion of hedges | $[0 - 1]$ | - |
| $\rho$ | Correlation between crops and hedges GRFs | 0.5 | - |
| **Parameters for population dynamic model** | | | |
| $D^v_2$ | 2D predator diffusion rate | $[0.0625 - 12]$ | $km^2d^{-1}$ |
| $m_v$ | Predator mortality rate | $[5 - 15]$ | $d^{-1}$ |
| $\beta$ | Predation rate | $[1 - 10]$ | $d^{-1}$ |
| $\rho_{21}$ | Predator migration rate from fields to hedges | 5 | - |
| $D^1_1$ | 1D predator diffusion rate | 12 | $km^2d^{-1}$ |
| $r_v$ | Predator intrinsic growth rate | $[10 - 20]$ | $d^{-1}$ |
| $K_{h_i}$ | Predator carrying capacity in hedge | 1 | - |
| $\rho_{12}$ | Predator migration rate from hedge to field | $[0 - 5]$ | - |
| $D^u_2$ | 2D pest diffusion rate | $[0.0625 - 12]$ | $km^2d^{-1}$ |
| $r_u$ | Pest intrinsic growth rate | $[10 - 20]$ | $d^{-1}$ |
| $C_{it}$ | Pest carrying capacity in crop fields | $C_{it} = \begin{cases} 20 & \text{no treatment} \\ 0.1 & \text{after the treatment} \end{cases}$ | - |
| $m_u$ | Pest mortality rate | $[5 - 15]$ | $d^{-1}$ |

### 4 Landscape discretisation for regression models

The spatial domain is discretised into small cells, different from the field polygons, and we assume a homogeneous point process intensity within each cell during each interval of time. Each combination of a spatial cell and a time unit represents a subset $A$ as landscape elementary unit, where we count the events occurring within it and evaluate additional information (*i.e.*, spatio-temporal covariates) associated to each cell. The discretisation is achieved through a triangulation mesh of space, whose construction puts focus on the influence of landscape structure on pest-predator dynamics. We take into account the patch structure of the landscape by differentiating between three types of cells: cells where exactly two patches are adjacent (located around the patch edge midpoints, with half of the edge length contained within the cell, representing movement corridors between exactly two neighbouring patches), cells where three patches are adjacent (around the vertices of the T-tessellation, containing 1/4 of each of the neigbhouring edge lengths, representing movement corridors around hubs where more than 2 patches come together), and cells at the center of each patch (representing areas that are separated away from edges and hedges, representing the patch inner core). This kind of discretisation allows us to focus on the influence of linear landscape elements represented by patch boundaries where hedges could be allocated, and we assume that the intensity function of the space-time point process is homogeneous within each cell for a given time step. Due to the reflecting boundary conditions of the population dynamics model, we observed a relatively large number of pest density peaks occurring at the boundaries of the study region, and we decided to remove all the boundary cells from the analysis to avoid an overly high influence of the boundary behaviour on the CBC analysis.

### 5 Predictor variables for pest dynamics

We evaluate spatio-temporal, spatial and population trait covariates for each cell of the mesh to relate the spatio-temporal event patterns, landscape structure and population dynamics. The full list of covariates is summarised in Table 2. Spatio-temporal covariates ($STC$) refer to the number of pest peaks, treatments and introductions that have occurred in previous time steps in the same cell or in neighbouring cells. This allows identifying spatio-temporal dynamics driven by preceding events at local scale. For example, we consider the number of treatments in each cell occurring at the previous time step to characterise recent spatio-temporal dynamics. We also evaluate the number of cumulated events up to two time steps before the present one to assess the influence of the cell's full local temporal dynamics in the past. Spatial covariates ($SC$) refer to geometrical features of the landscape patch tessellation and the land use allocation to evaluate the effect of the configuration and composition of the landscape. For example, to evaluate landscape configuration properties, we consider a three-level categorical variable to indicate if the cell connects exactly two, or more than two patches, to assess its spatial position that could be important to assess fluxes among patches. We evaluate the landscape composition at local scale through a variable measuring the percentage of crop and hedge within a buffer centered on the centroid of each cell, and at global scale through a variable measuring the percentage of crop and hedge in the whole landscape. This enables an assessment of how the crop extent and the hedge network may locally and globally influence the population dynamics leading to the number of events within each cell. Specifically, since the buffer aims to characterise variables at local scale, the buffer diameter is adapted to pest growth rate $Rprey$ and dispersal $Dprey$, which determine the spreading speed of the population. The spreading speed can be evaluated as $v = 2\sqrt{Rprey * Dprey}$. Then, the travelled space (*i.e.*, the front of the population) is computed by the classical formulation $s = v \times t$, where $s$ is the travelled space, identifying the front position, and $t$ is the time, set equal to 0.01, as in the population dynamics model. Among all the possible simulated configurations of $Rprey$ and $Dprey$, resulting in highly different buffer sizes $s$, we group configurations in three classes defined by the $10\%-, 50\%-, 90\%-$percentile of the population front $s$. Then, the buffer diameter size is set equal to the population front $s$ used as an approximation of the local range where the population could move during a time step starting from the center of a cell. The three resulting buffers have a diameter equal to 0.09, 0.19 and 0.26 $Km$ for the $10\%-, 50\%-, 90\%-$percentile of $s$, respectively. To each pair of $Rprey$ and $Dprey$ is assigned the corresponding buffer, where we then calculate covariates of crop and hedge proportion. For population dynamics covariates ($PDC$), we select variables related to individual mobility, such as species dispersal in fields and spillover from hedge to fields. We select these species trait variables as they are directly related to the species' capability to move within the landscape.

Table 2: List of covariates of regression models.

| Ref. ID | Covariate name | Spatial reference | Range | Unit | Description |
|---|---|---|---|---|---|
| **Spatio-temporal** ($STC$) | | | | | |
| 1 | tr_patch($t-1$) | patch | 0-40 | - | No. of treatments in the patch at $t-1$ |
| 2 | tr_patch_cum($t-2$) | patch | 0-97 | - | No. of treatments in the patch cumulated up to $t-2$ |
| 3 | tr_Nb.patch($t-1$) | patch | 0-337 | - | No. of treatments in neighbor patches at $t-1$ |
| 4 | tr_Nb.patch_cum($t-2$) | patch | 0-861 | - | No. of treatments in neighbor patches cumulated up to $t-2$ |
| 5 | pk_cell($t-1$) | cell | 0-15 | - | No. of pest density peaks at $t-1$ |
| 6 | pk_cell_cum($t-2$) | cell | 0-36 | - | No. of pest density peaks cumulated up to $t-2$ |
| 7 | pk_Nb.cell($t-1$) | cell | 0-45 | - | No. of pest density peaks in neighbor cells at $t-1$ |
| 8 | pk_Nb.cell_cum($t-2$) | cell | 0-97 | - | No. of pest density peaks in neighbor cells cumulated up to $t-2$ |
| 9 | int_cell($t-1$) | cell | 0-30 | - | No. of pest introduction in cell at $t-1$ |
| 10 | int_cell_cum($t-2$) | cell | 0-30 | - | No. of pest introduction in cell cumulated up to $t-2$ |
| 11 | int_Nb.cell($t-1$) | cell | 0-30 | - | No. of pest introduction in neighbor cells at $t-1$ |
| 12 | int_Nb.cell_cum($t-2$) | cell | 0-39 | - | No. of pest introduction in neighbor cells cumulated up $t-2$ |
| **Spatial** ($SC$) | | | | | |
| 13 | Area | cell | 0-0.069 | $km^2$ | Cell dimension |
| 14 | cell_2patch | cell | 0-1 | - | Binary, it is = 1 if the cell is among 2 patches, 0 otherwise |
| 15 | cell_3patch | cell | 0-1 | - | Binary, it is = 1 if the cell is among 3 or more patches, 0 otherwise |
| 16 | %H_Buffer | buffer | 0-1 | % | Percentage of hedges within the buffer centered in the cell |
| 17 | %C_Buffer | buffer | 0-1 | % | Percentage of crops within the buffer centered in the cell |
| 18 | Aggr | landscape | 0-5.54 | - | Landscape crop and hedge aggregation |
| 19 | %C_Land | landscape | 0-1 | % | Landscape crop proportion |
| 20 | %H_Land | landscape | 0-1 | % | Landscape hedge proportion |
| **Population dynamics** ($PDC$) | | | | | |
| 21 | Dprey | landscape | 0.06-12 | $km^2d^{-1}$ | Pest diffusion in crop patch |
| 22 | Dpred | landscape | 0.07-12 | $km^2d^{-1}$ | Predator diffusion in crop patch |
| 23 | D12pred | landscape | 0.1-1 | | Predator diffusion from hedge to crop |

### 6 Correlation analysis of covariates

We assessed the correlations among the selected covariates defined for each cell of the domain and for each time step Figure 2. The main positive correlations arise for the covariate % of culture and % of hedge within the buffer, and the corresponding % of culture and hedge in the whole landscape, respectively. Other strong positive correlations are shown by the covariate evaluating the number of peaks and treatments at the previous time step or cumulated up to $(t-2)$ in the same cell or in the neighbouring ones. This highlights that locations favourable to high pest density will always experience pest outbreaks even after treatments.

### 7 Residual analysis of regression models

A residual analysis is performed to evaluate if the predicted values obtained by the GLMs are homogeneously distributed in tspace and time for the models of intensity peak occurrence and of peak value of pest density. Residuals are computed as the difference among the predicted values and the observed spatially explicit population model outcomes.

We analyse the residuals to check the residual homogeneity over space and the time (Figures 3, 4 and 5). We can conclude that the models defined for pest peak number and value are able to capture the variability of observed data (*i.e.*, population dynamic model outputs) in time and space (Figures 3) without any systematic biases. We can recognise the already discussed spatial trend depending on cell position: the first 200 cells are cells intersecting 3 patches, the cells among 200 and 600 are center cells. As to the model for peak number, higher intensity takes place in cells located over patch boundaries, while in the model for pest peak density value higher values take place in center cells. The temporal dynamics are very well captured and show a slow increasing trend for pest outbreak occurrence up to a maximum value at last time step, and a fast and drastic increasing trend for pest density values with respect to time. In Figure 4 and 5 a) and d), we can see that the residuals are relatively small and almost homogeneously distributed for both models in space (a) and time (d). This can be also verified in the map of the spatial domain discretised in cells where we visualise the error within each cell. Here, we remark that for the model of peak numbers, larger errors occur in cells intersecting 2 or more patches, where the intensity of occurrence is slightly overestimated. By contrast, in the model of peak valuse, larger errors occur in the relatively large cells centred in patches, where the pest density concentration tends to be slightly underestimated. In Figure 4 and 5 e), we report

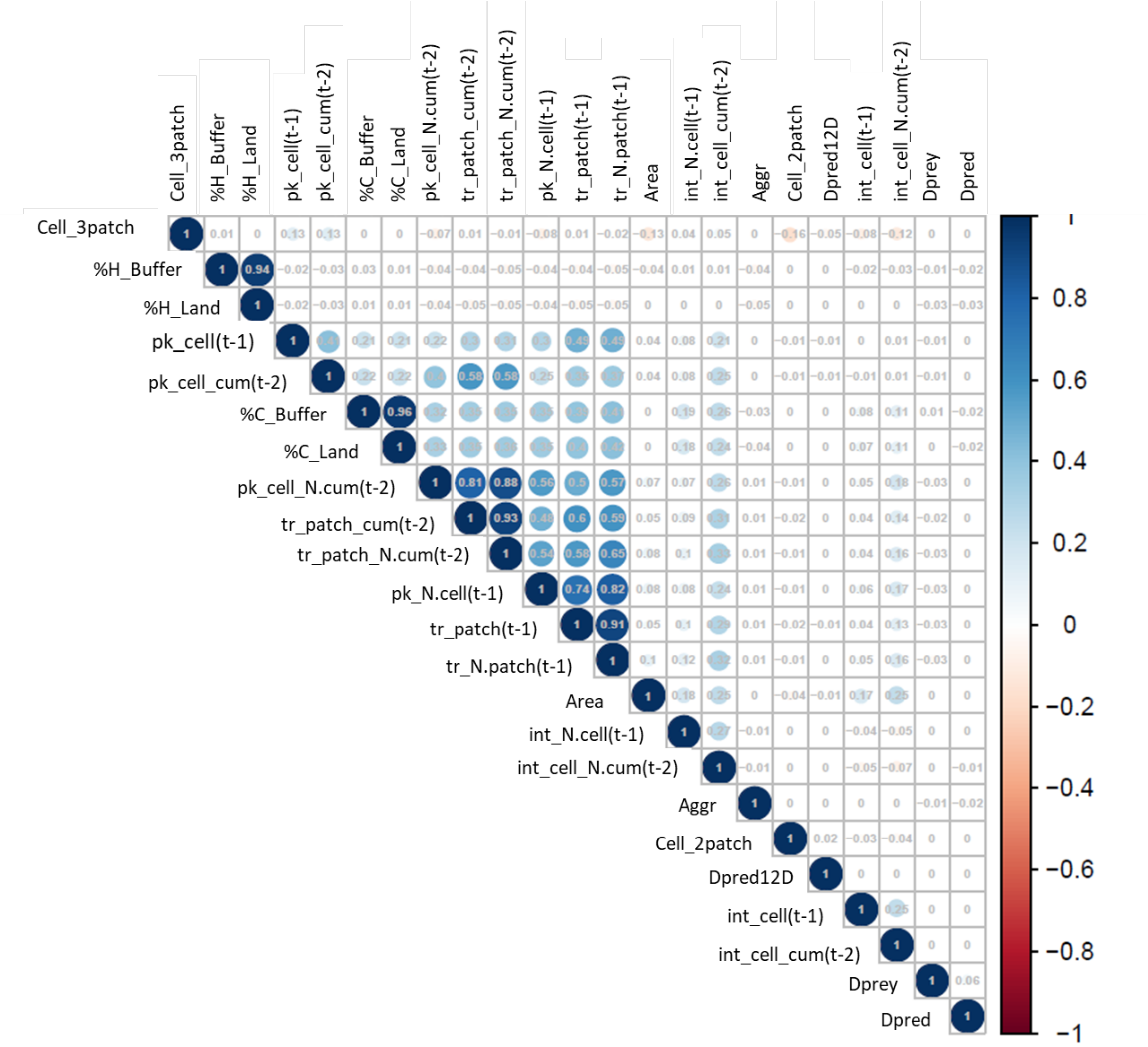


Figure 2: Empirical Pearson correlations among covariates selected for the occurrence intensity of pest density peaks and for the model for the peak value of the pest density.

the same plot as in Figures 3a but for each time step, in order to verify that also the dynamical behaviour is well captured for each time step, and not only on the temporal average. Lastly, the spatial variograms of residuals, 4 and 5 panels c), confirm that there is the same variability among cells over the distance, *i.e.*, there is no spatial structure left in the residuals.

Finally, we show the estimated temporal effect (*i.e.*, the intercept specific to each time interval) in the regression models for the occurrence intensity of pest density peaks (Figure 6a) and for the peak value of pest density (Figure 6b). We had introduced this temporal effect in order to better take into account and explain the temporal dynamics of the population, which were not fully explained by the other covariates. As for the occurrence intensity of pest density peaks (Figure 6a), the time-varying intercept is positive and it increases with time up to a constant value. This reflects that the number of pest outbreaks increases through time as the pest population grows and colonises the spatial domain. As for the peak value of pest density (Figure 6b), the time-varying intercept is negative and shows a relatively weak amplitude. It decreases with time, such that the maximum value of pest outbreaks occurs in the first time steps, probably when clusters of pest concentration arise that are not well homogenised over the spatial domain. Afterwards, when the pest population expands, there is a dilution effect that leads to a decrease in pest density peak values.

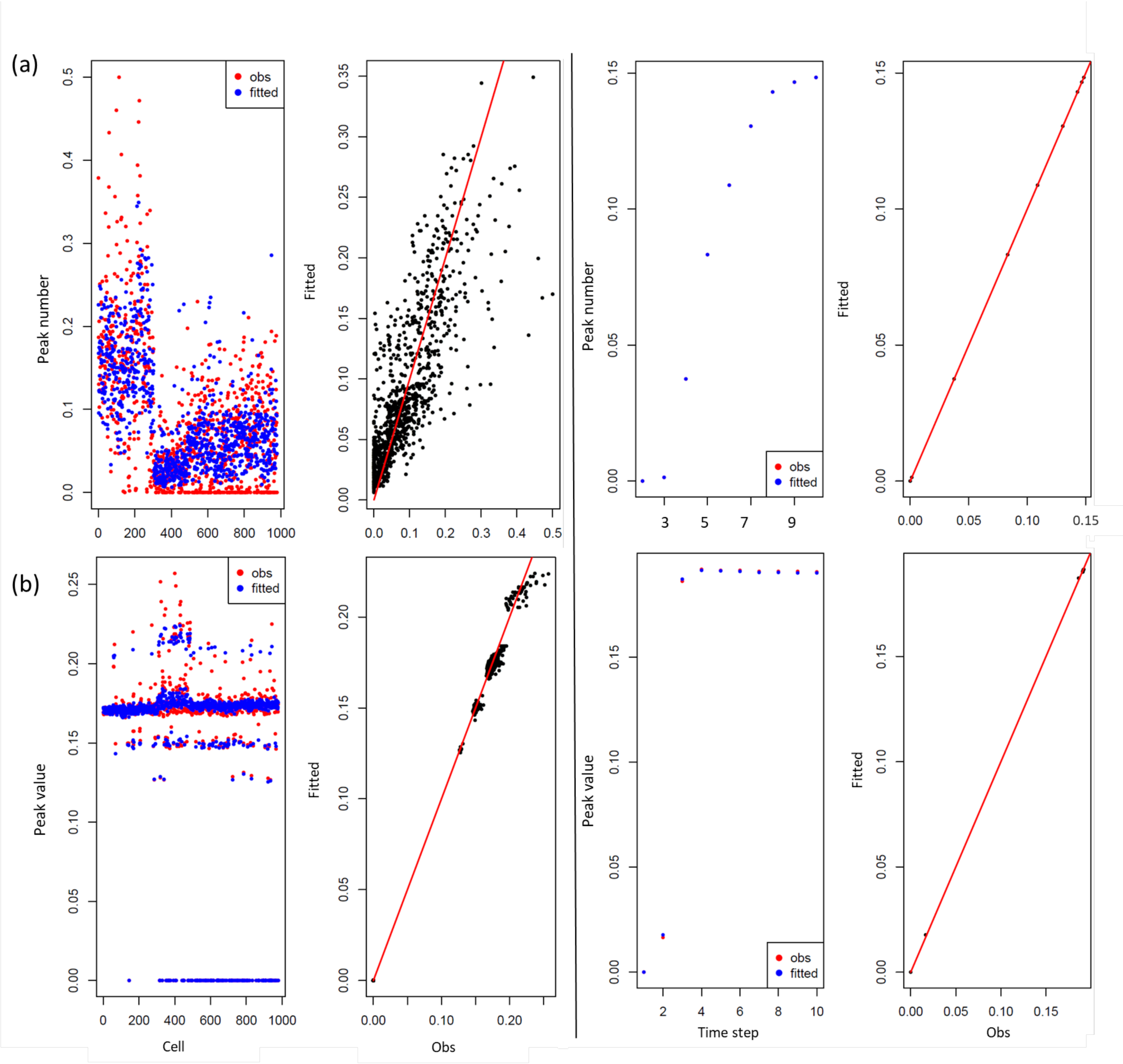


Figure 3: Residual analysis showing observations, *i.e.*, model output (red dots), with predicted (blue dots) occurrence of peaks (Panel a) and of peak maximum values (Panel b) over space (left panels, by averaging over time) and over time (right panels, by averaging over space).

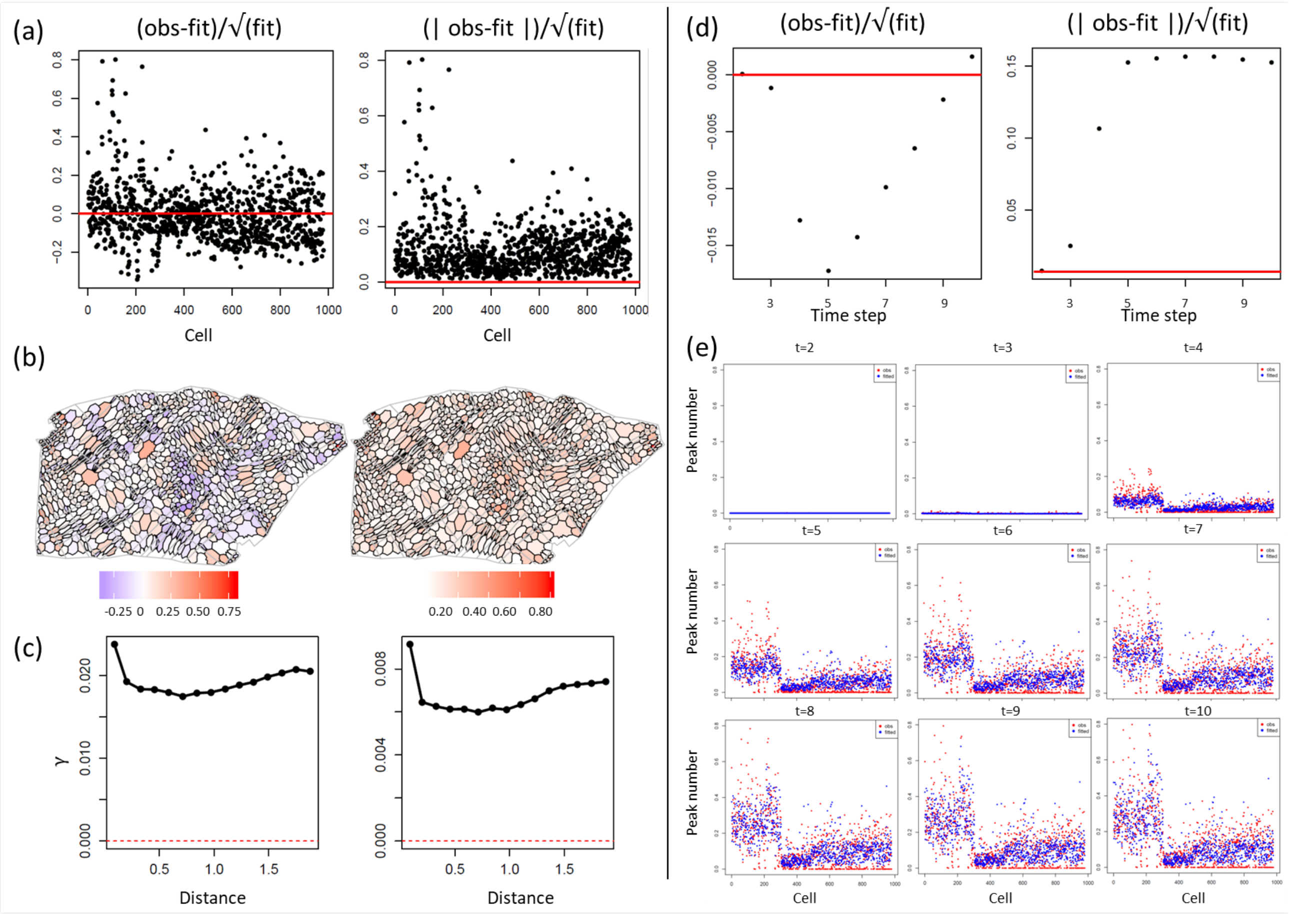


Figure 4: Residual analysis for the model of the occurrence intensity of peaks. Left: Spatial residual analysis. Panel a) Residuals scaled over the fitted values for each cell averaged over time steps; Panel b) Visualisation of the scaled residuals for each cell averaged over time steps; Panel c) Variogram of the scaled residuals. Right: Temporal analysis. Panel d) Residuals scaled over the fitted values for each time step averaged over cells; Panel e) Observations (red dots) and fitted values (blue values) for each time step.

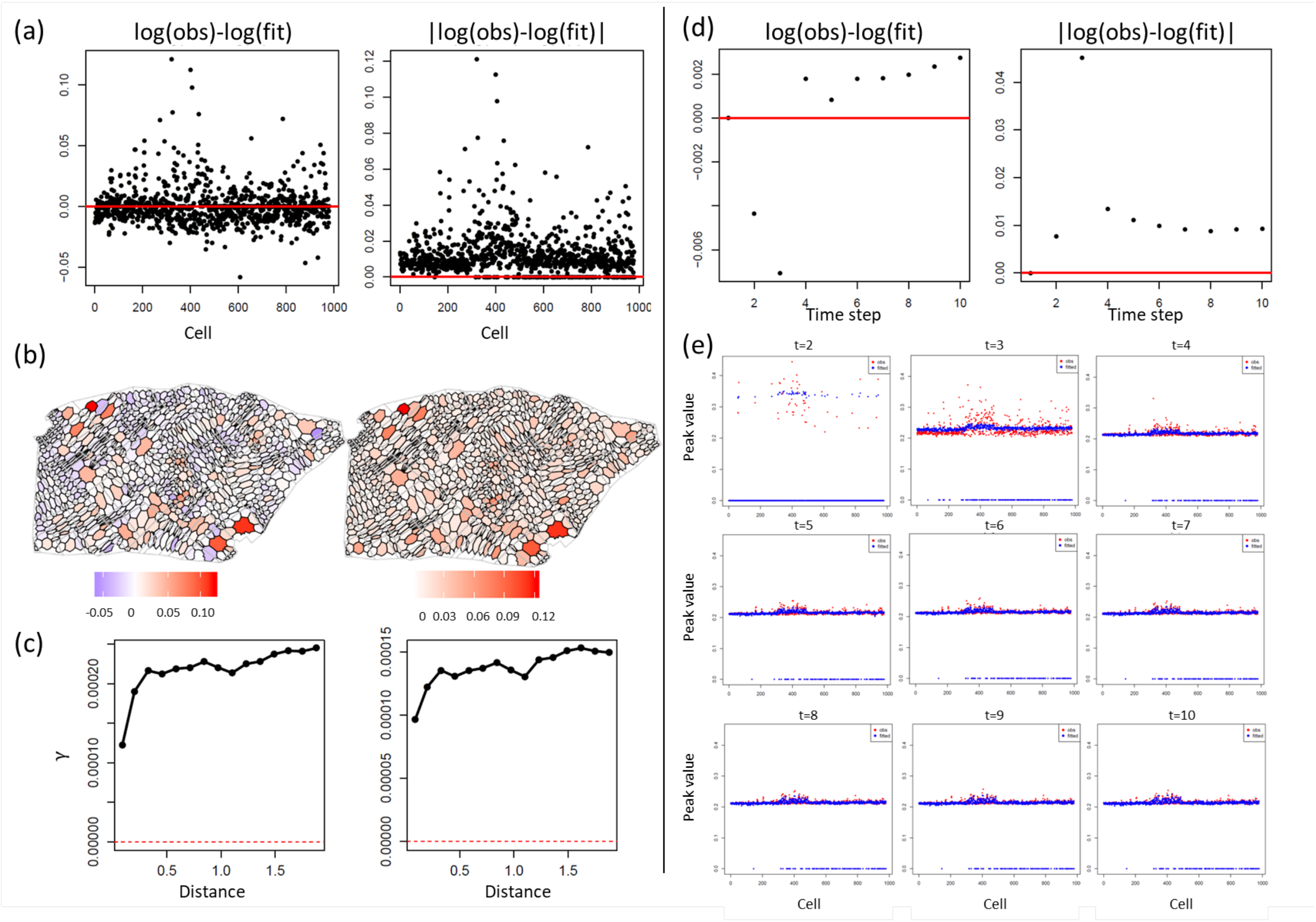


Figure 5: Residual analysis for the model of the peak values. The results presented follow the same structure as in Figure 4

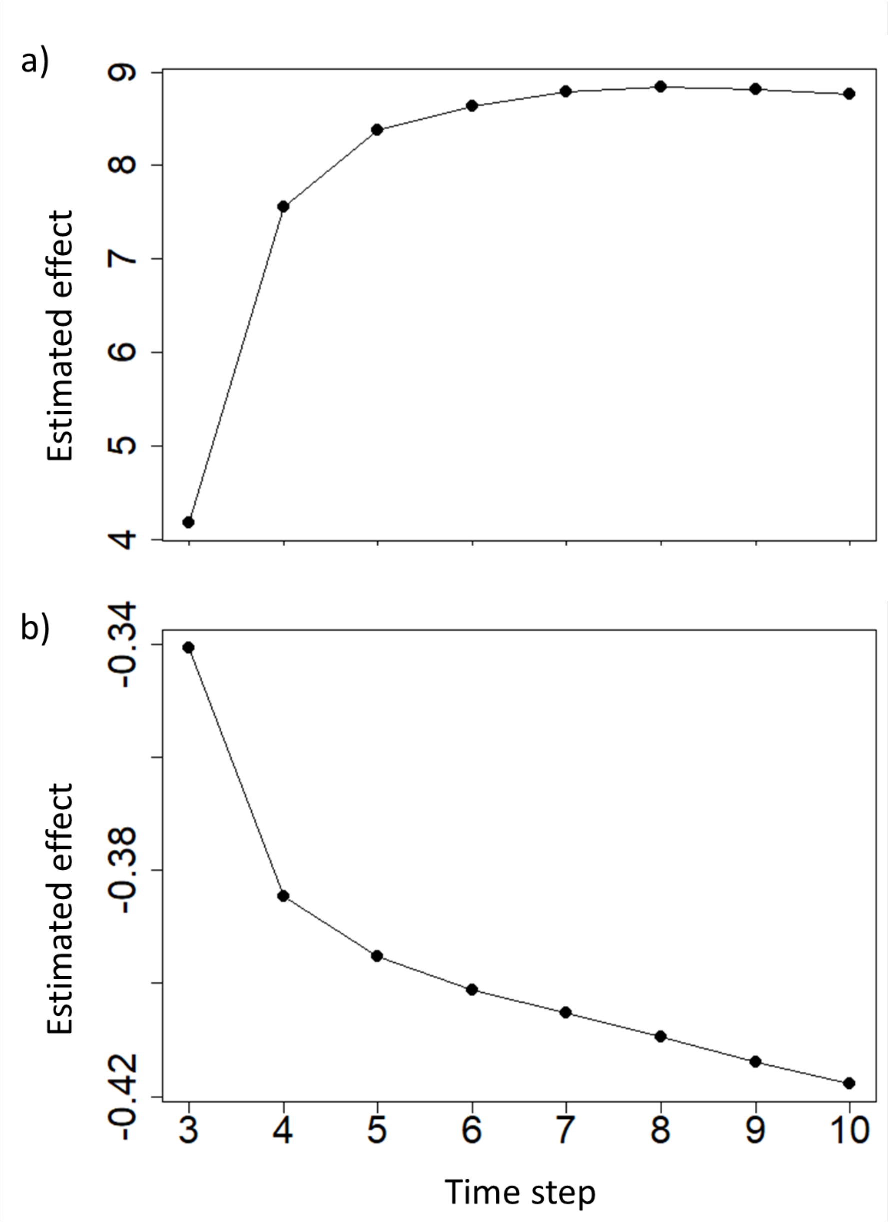


Figure 6: Estimated temporal effects for the occurrence intensity of pest density peaks (Panel a) and for the peak values of pest density (Panel b).
